# The ESX-1 secretion system senses bacterial contact and prepares mycobacteria for environmental adaptation

**DOI:** 10.1101/2021.10.17.464699

**Authors:** Nadia Herrera, Pascal D. Odermatt, Mark Voorhies, Rachel Nakagawa, Anita Sil, Fred Chang, Oren S. Rosenberg

**Affiliations:** Department of Medicine, University of California San Francisco, San Francisco, California, USA; Department of Biochemistry and Biophysics, University of California San Francisco, San Francisco, California, USA; Chan Zuckerberg Biohub, San Francisco, California, USA; Department of Cell and Tissue Biology, University of California San Francisco, San Francisco, California, USA; Department of Microbiology and Immunology, University of California San Francisco, San Francisco, California, USA

## Abstract

The ESX-1 system (6-kDa early secretory antigenic target (ESAT-6) secretion system-1) is essential for *Mycobacterium tuberculosis* pathogenesis and conjugal transfer in *Mycobacterium smegmatis*, yet little is known about how its function is regulated. Live-cell fluorescence microscopy showed natively expressed ESX-1 was organized into distinct foci predominantly observed at cell-cell contacts. These foci formed when two cells touched and required a fully assembled ESX-1 system in both bacteria, suggesting the generation of an ESX-1 megacomplex across multiple membranes. The emergence of ESX-1 foci and ESX-1 secretion was environmentally dependent: foci formed in low nitrogen environments in which secretion was suppressed, yet with increasing concentrations of nitrogen, ESX-1 systems diffused along the plasma membrane and secretion was activated. Genome-wide transcriptional profiling revealed ESX-1 dependent induction of genes required for the SOS response and error prone DNA replication in high nitrogen. Based on these findings, we propose a new model of ESX-1 function where ESX-1 localization and secretion are responsive to nitrogen levels and form an integral node in the mycobacterial response to neighboring cells and environmental adaptation.

## Introduction

Mycobacteria utilize ESX (6-kDa early secretory antigenic target (ESAT-6) secretion) systems to shuttle specialized substrates across a diderm cell wall. ESX-like systems are widely conserved in saprophytic bacteria, including the actinobacteria and firmicute phyla, but they have greatly expanded in the mycobacteria (Baptista et al., 2013; Burts et al., 2008, 2005; Dumas et al., 2016; Garufi et al., 2008; Gey Van Pittius et al., 2001; Huppert et al., 2014; Way and Wilson, 2005). There are five paralogous ESX systems in mycobacteria, termed ESX-1 – ESX-5, that stem from the ancestral ESX-4 which is most closely related to other systems in the broader phyla (Newton-Foot et al., 2016). The ESX systems have been implicated in the core characteristics of mycobacteria including virulence (ESX-1) (Cole et al., 1998; Sørensen et al., 1995), metal homeostasis (ESX-3) (Serafini et al., 2013, 2009, Siegrist et al., 2014, 2009; Tufariello et al., 2016), and phosphate regulation(ESX-5) (Elliott and Tischler, 2016). Single particle cryo-electron microscopy showed that two ESX systems (ESX-3 and ESX-5) share a defined structure, suggesting that the assembled ESX systems likely share an underlying biochemical mechanism (Beckham et al., 2021, 2017; Famelis et al., 2019; Poweleit et al., 2019), but have different biological functions possibly based in the use of different substrates or accessory factors (Beckham et al., 2017; Famelis et al., 2019; Phan et al., 2018; Poweleit et al., 2019; Siegrist et al., 2014). ESX loci in mycobacteria consist of the ESX conserved components (Ecc’s), which include membrane-embedded components (EccB, EccC, EccD, and EccE), secretion substrates such as the Esx, PE, PPE, Esp proteins, motor ATPases such as EccCb and EccA, and regulatory elements (Figure 1A) (Berthet et al., 1998; Pallen, 2002).

**Figure 1:**
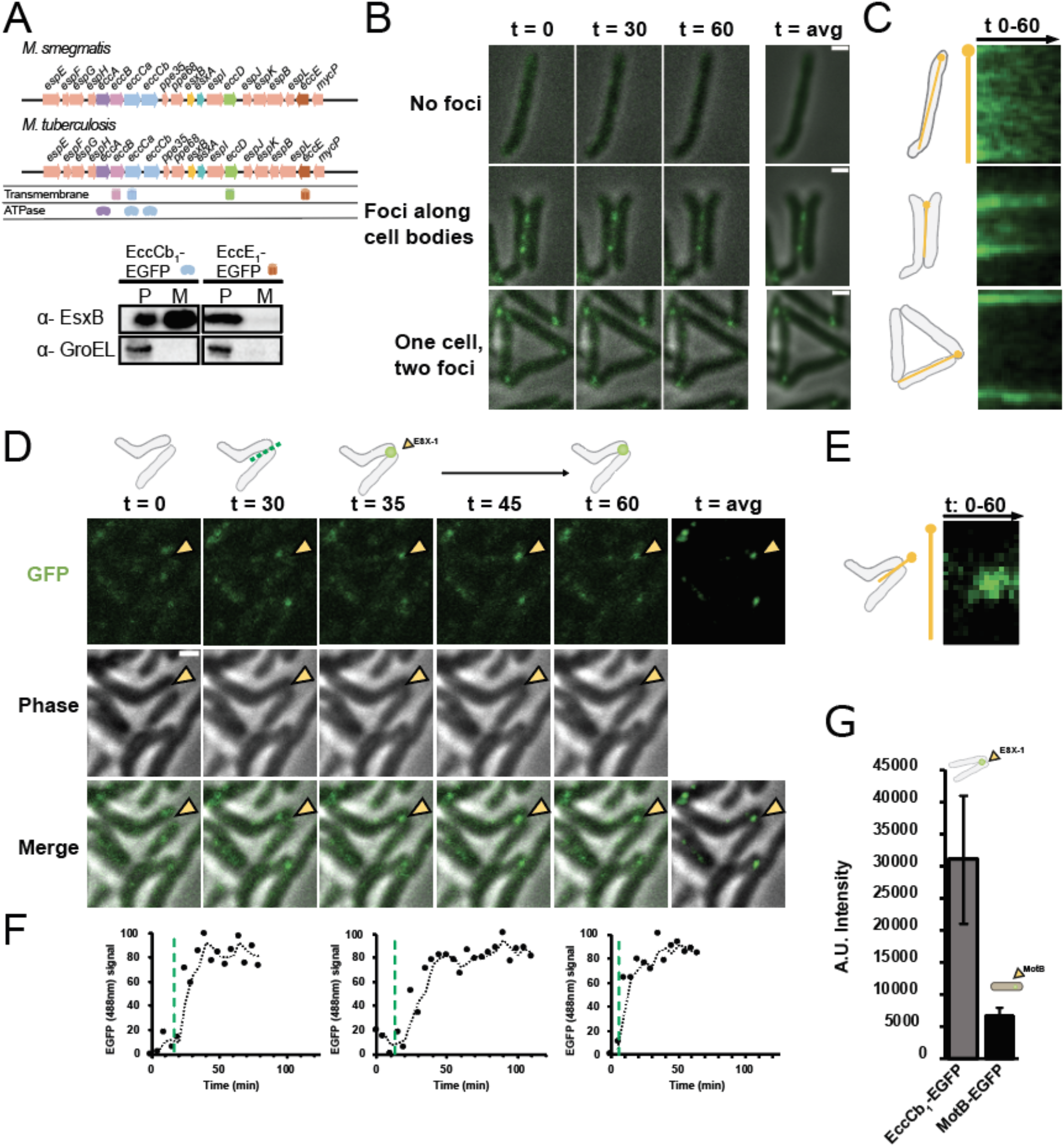
ESX-1 forms foci at cell-cell contacts. A) Top-Organization of ESX-1 operon in *M. smegmatis* and *M. tuberculosis*. Proteins with transmembrane domains and ATPase cassettes are defined. Bottom-Secretion assay in EccCb_1_-EGFP and EccE_1_-EGFP strains of *M. smegmatis*. Schematic of gene within the operon is shown to the right of the label. Western blot shows the ESX-1 substrate EsxB secreted into the medium. P represents pellets and M represents medium, and anti-GroEL antibody shows the integrity of the loaded samples. B) Confocal microscopy images of distinct cell-cell contacts at time points 0, 30, and 60 minutes; only merged image depicted. Time averaged (t avg.) images represent averaged images over entire time course. C) Schema of images in A. The yellow pin depicts location of kymograph of GFP signal through the time-lapse acquisition in panel A. D) Top-Schema illustrates a cartoon rendition of EGFP foci formation event. Bottom-Confocal microscopy images of growing cultures at time points 0, 30 and 35 minutes, and average, an average of all time points. Imaging channels include GFP, phase, and merged channels of two cells encountering each other. Point of cell-cell contact depicted by yellow arrowhead. E) Kymograph represents EGFP signal throughout duration of the experiment at the cell-cell contact site. F) Normalized integrated fluorescence measurements of 3 distinct cell-cell contact events. Green dashed line indicates time of contact as determined by phase images. Top graph corresponds to images in 1A. G) Fluorescence intensity measurements of EccCb_1_-EGFP in both *M. smegmatis* (grey, N = 53) and MotB-EGFP *E. coli* (black, N = 101). Foci were quantified for three biological replicates. Error bars indicate standard deviation of measurements. Scale bar located on top right corner of images, 1µm.

ESX-1, the first described ESX system, was identified through genomic studies as the key genetic difference between *Mycobacterium tuberculosis* and the attenuated *Mycobacterium bovis* BCG vaccine strain (Hsu et al., 2003; Pym et al., 2003, 2002; Sassetti and Rubin, 2003; Stanley et al., 2003). The ESX-1 system is highly conserved in the non-pathogenic model organism, *Mycobacterium smegmatis* whose ESX-1 system shares a 72% nucleotide sequence conservation with *M. tuberculosis* ESX-1 across the protein coding regions of ESX-1 components (Converse and Cox, 2005). As *M. smegmatis* is non-pathogenic, ESX-1 clearly has additional functions besides those associated with pathogenesis. For instance, it has been associated with regulation of conjugal DNA transfer in the *M. smegmatis* strain MC^2^155 and other mycobacterial species (Gray et al., 2016; Gröschel et al., 2016). However, in general, the functions of ESX-1 secretion remain poorly understood.

The localization of ESX-1 in bacterial cells remains unclear. Previous studies localized overexpressed GFP-fusion of ESX-1 associated proteins EccCb_1_ and EccE_1_ to the polar regions of *M. smegmatis*, and *M. tuberculosis* (Soler-Arnedo et al., 2020; Wirth et al., 2012). Polar localization was also seen in *M. marinum* using immunofluorescence on a cell wall deficient mutant (Δ*kasB*) (Carlsson et al., 2009). Combined, protein overexpression and cell wall interruption may disturb the physiological localization pattern of a membrane complex, leaving the localization of natively expressed ESX-1 components as an open question. Furthermore, there is growing evidence that ESX-1 secretion systems are regulated by environmental factors during *M. tuberculosis* infection (Berthet et al., 1998; Fortune et al., 2005). In addition, in *M. smegmatis*, ESX-1 secretion was found to be active when cells were grown on Sauton’s medium, and largely inactive in 7H9 medium (Converse and Cox, 2005). The ramifications of ESX-1 regulation are yet to be explored.

In this work we constructed functional GFP fusions expressed at endogenous levels to study the localization of ESX-1 components. We found that ESX-1 formed discrete foci at either side of a cell-cell contact in cells grown in 7H9 medium, a condition in which ESX-1-mediated EsxB secretion was inhibited. Conversely, ESX-1 was localized diffusely around the membrane in Sauton’s medium, when ESX-1 secretion of EsxB was active. We show that the increase in nitrogen levels in Sauton’s medium was sufficient to induce both re-localization and activation of ESX-1 secretion of EsxB. We used RNAseq to probe the physiological function of ESX-1 and discovered that ESX-1 was necessary for activating the mycobacterial SOS response to nitrogen addition. Taken together, these findings, suggested an unexpected function of the ESX-1 secretion system in regulating stress responses in high nitrogen environments that may inform on its role in mycobacterial pathogenesis.

## Results

### ESX-1 forms stable foci at cell-cell contacts

Prior studies reported the localization of heterologous ESX-1 components upon overexpression of plasmid-based EGFP fusions (Soler-Arnedo et al., 2020; Wirth et al., 2012), which in some cases, leads to non-physiological localization. To investigate the localization of native ESX-1 expressed at endogenous levels, we introduced an EGFP tag into multiple ESX-1 components in the chromosome. We determined that EccCb_1_-EGFP was a functional EGFP fusion, as shown by a secretion assay probing for EsxB in the culture medium although other fusions were not functional (Figure 1A). We grew these cells to exponential phase in 7H9 liquid medium and then mounted them into microfluidic chambers for time lapse spinning disc confocal microscopy.

In the minority of cells that were not physically contacting another one, EccCb_1_-EGFP exhibited a dim localization around the whole plasma membrane (Figure 1B, top panel). This localization was accentuated by time averaging the images (t avg.). Kymography analysis showed that this intensity was maintained over time (Figure 1C, top). However, most cells clumped together in large aggregates in 7H9. In cells that were contacting others, we found that ESX-1 components formed discrete foci at regions of cell-cell contact (Figure 1D middle and bottom panels). These foci were observed at the contact site between cells that occurred either along the cell body (Figure 1B middle panel) or between two cell poles (Figure 1D bottom panel). The foci were stable in intensity and were immobile for the duration of the measurement (Figure 1D, t = 0 min – t = 60 min and t avg.). The stability of the foci is illustrated in a kymograph (Figure 1C middle and bottom panels). This observation demonstrates that ESX-1 foci form at cell-cell contact sites and are not limited to cell poles. We also confirmed that EccCb_1_-EGFP plasmid-based overexpression caused foci formation at the poles, regardless of cell-cell contact (Supplemental Figure 1) suggesting that previously reported polar localization may be due to overexpression of and subsequent EGFP self-interaction (Landgraf et al., 2012). In addition, we show that endogenous expression of the monomeric construct of EGFP, mEGFPmut3, yields similar observations of foci at cell-cell junctions and does not differ from our EGFP observations (Supplemental Figure 2).

We used time-lapse microscopy to investigate the dynamics of focus formation (Figure 1D). Growing cells were observed while contacting each other and forming an ESX-1 focus on the cell-cell contact point. A representative example is shown in Figure 1C. At t = 0 min when the cells were near but not touching each other, there were no detectable foci. At t = 30 min multiple dim foci appeared along the contact region. By t = 35 min onward there was a single, persistent ESX-1 focus on the contact site that persisted. A kymograph of the entire time-lapse acquisition at the cell-cell contact site illustrates this behavior over time (Figure 1E). EGFP intensity plots show that the foci form within 5 minutes (Figure 1F) in a few examples. These images provide a striking demonstration that ESX-1 focus formation accompanies cell-cell contact.

### ESX-1 foci form large oligomers at cell-cell contact sites

The discrete ESX-1 foci suggested that ESX-1 formed large oligomeric assemblies at the membrane at cell-cell contacts. To quantify the number of ESX-1 complexes at these sites we compared the fluorescent intensity of EccCb_1_-EGFP foci to the MotB-EGFP complex which has been reported to contain 22 +/-4 EGFP molecules in each focus (Coffman and Wu, 2012; Leake et al., 2006; Pan et al., 2014) (Supplemental Figure 3). Measurements of EGFP intensity of these foci indicated that the EccCb_1_-EGFP foci were 6-fold more intense than MotB-EGFP foci (Figure 1G). Measured intensities were uniformly distributed, following a gaussian distribution. This analysis suggests that a single ESX-1 focus contains approximately 132 individual EGFP molecules. Considering the predicted hexameric structure of ESX systems (Famelis et al., 2019; Poweleit et al., 2019), this suggests roughly 22 hexameric complexes of ESX-1 at each focus. This arrangement suggests that ESX-1 forms a large complex at the membrane of cell-cell contacts.

### ESX-1 foci formation requires an intact ESX-1 system

We next sought to address what molecular components of the ESX-1 system are needed for focus formation. EccCb_1_ was endogenously labeled with EGFP in strains with a series of gene deletions in the ESX-1 operon. In our analysis (Figure 2) we scored whether a focus was present at cell-cell contacts (grey bars, depicted by schema on top right of graph). Deletion of *eccB*_*1*_ resulted in decreased percentage of cells exhibiting EccCb_1_-EGFP foci at cell-cell contacts with only 30% of cell contacts displaying foci, while *eccD*_*1*_ and *eccE*_*1*_ deletions resulted in a complete lack of foci at cell contacts. In comparison, wildtype cells formed foci at all cell contacts 100% of the time (Figure 2A, B). Interestingly, upon deletion of *esxB*, a major secreted product of ESX-1, focus formation remains largely unaffected, with 99% of cell contacts displaying foci (Figure 2A, bottom panel and Figure 2B). We complemented the deleted *ecc* components in the appropriate strains using an integrative plasmid harboring the gene of interest under a neutral promoter. Analysis of cell-cell contacts demonstrated that focus formation was largely rescued (Figure 2B). These results indicated that ESX-1 focus formation was dependent on integral membrane components within the Ecc’s of ESX1 but did not require its substrate, EsxB, or the secretion of the substrates dependent on EsxB.

**Figure 2:**
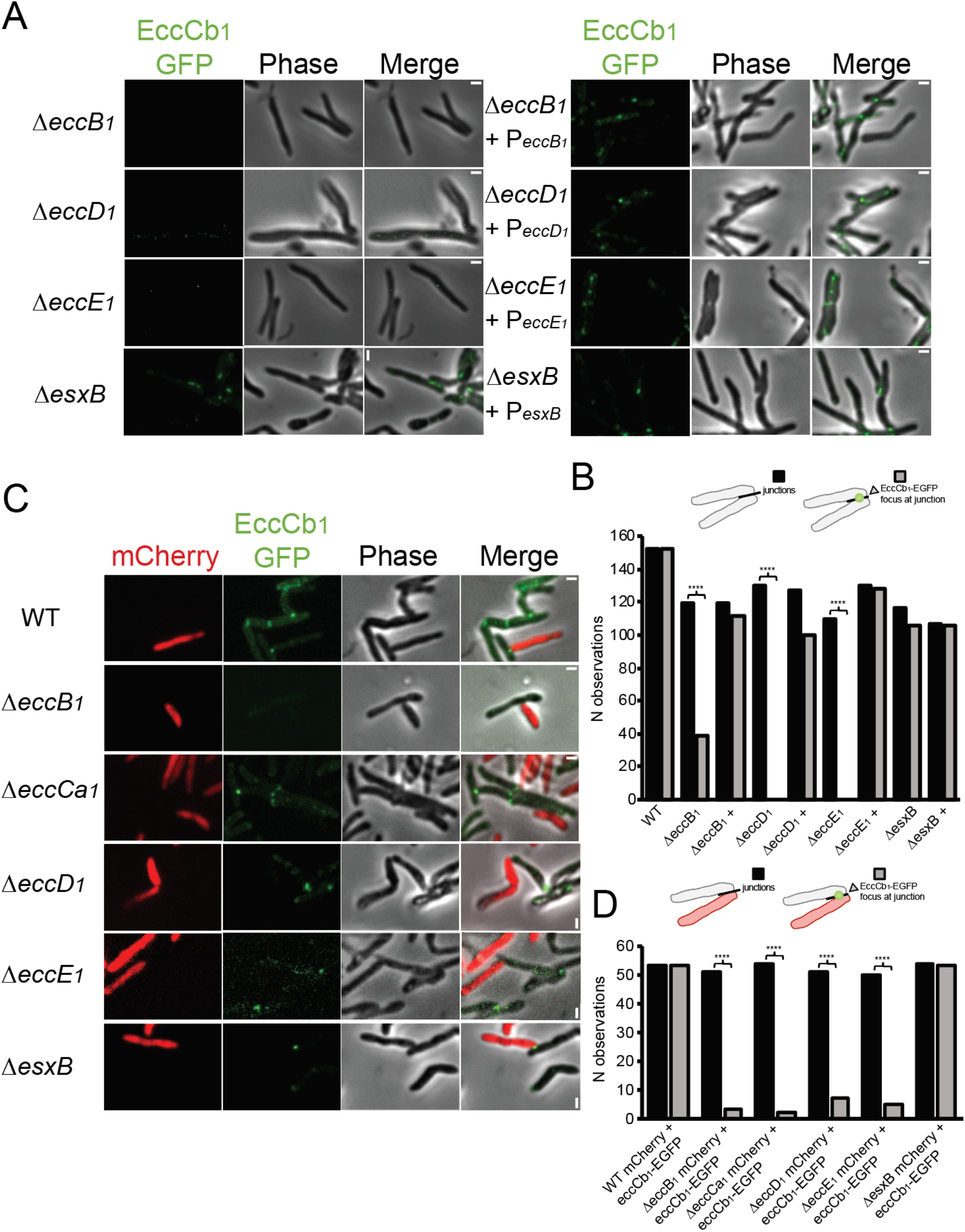
ESX-1 is required at both interfaces to form a focus. Time averaged confocal images acquired every 5 minutes for the duration of an hour are depicted. A) ESX-1 essential conserved components (ecc’s) were knocked out and EccCb1-EGFP was used as a tracer. Left panel shows GFP, phase and merged channels. Blue arrowheads indicate contacts with no foci at cell-cell contacts. Right panel shows GFP, phase and merged channels, yellow arrowheads represent cell-cell contacts where foci formation was restored upon complementation of the knocked out ecc component. B) Foci were quantified for three biological replicates, across 100+ contacts per strain (reported on Y-axis as N observations) in both knockouts of ESX-1 and complement strains. Difference between both measurements is statistically significant per student’s t-test, **** P < 0.0001. C) Co-cultures of EccCb_1_-EGFP with wildtype, Δ*eccB*_*1*_, Δ*eccCa*_*1*_, Δ*eccD*_*1*_, Δ*eccE*_*1*_, Δ*esxB* (top to expressing mCherry are shown (top to bottom). Schema on the left represents the strains captured in the images. Foci at cell-cell contacts are outlined by yellow arrowheads, while cell-cell contacts lacking foci are outlined by blue arrowheads. Scale bar located on top right corner of images, 1µm. D) Foci were quantified for three biological replicates, across 50+ contacts per strain across the panel of co-cultures (reported on Y-axis as N observations). Difference between both measurements is statistically significant per student’s t-test, **** P < 0.0001. Scale bar located on top right corner of images, 1µm.

### ESX-1 forms a megacomplex across two contacting cells

We determined whether ESX-1 complexes at cell-cell contacts were a one or two-sided interaction. To address this question, we used a co-culturing experiment in which strains expressing EccCb_1_-EGFP were mixed with *ecc* knockout strains marked with a cytoplasmic mCherry and assayed whether EccCb_1_-EGFP foci were detected at cell-cell contact sites. As shown by representative time averaged images, focus formation was induced between wild-type and Δ*esxB* strains, whereas all *ecc* knockout strains were largely unable to induce foci formation in the other cell (Figure 2C). Focus formation in P_mCherry_ wild-type and Δ*esxB* cells retained ∼ 99% focus formation at cell-cell contacts with EccCb_1_-EGFP cells (Figure 2D). Focus formation at cell-cell contacts between P_mCherry_ -ecc knockouts and EccCb_1_-EGFP labeled cells was as follows: in Δ*eccB*_*1*_-P_mCherry_ 5%, Δ*eccCa*_*1*_-P_mCherry_ 3%, Δ*eccD*_*1*_-P_mCherry_ 13%, and Δ*eccE*_*1*_-P_mCherry_ 10% (Figure 2D). In all instances, foci still formed between EccCb_1_-EGFP labeled cells, indicating that focus formation was unaffected by the co-culture milieu (Figure 2C). We concluded ESX-1 focus formation required assembly of the ESX-1 complex in both contact cells, suggesting that ESX-1 clusters at the cell-cell contact sites on both plasma membranes, stabilizing each other, and forming a megacomplex that includes across both cell membranes

### High nitrogen concentrations in growth medium triggers ESX-1 secretion in *M. smegmatis*

We next investigated if focus formation was dependent on environmental conditions. Observations of ESX-1 focus formation were made in 7H9 medium, however, previous studies (Converse and Cox, 2005) showed that cultures grown in 7H9 medium suppressed ESX-1 secretion of EsxB, while those grown in Sauton’s media were proficient in EsxB secretion (reproduced in Figure 3A). We tested whether these differences in medium affected ESX-1 focus formation. Cells were grown to exponential phase in either 7H9 or Sauton’s medium and mounted into microfluidic chambers for time lapse spinning disc confocal microscopy. Time-averaged images of cells grown in secretion-inducing Sauton’s medium (Figure 3D and 3E) revealed the absence of ESX-1 foci at cell-cell junctions, while time averaged images of cells grown in 7H9 exhibited ESX-1 foci. In Sauton’s medium the EGFP signal was distributed throughout the plasma membrane, regardless of contact site with surrounding cells (Figure 3E).

**Figure 3:**
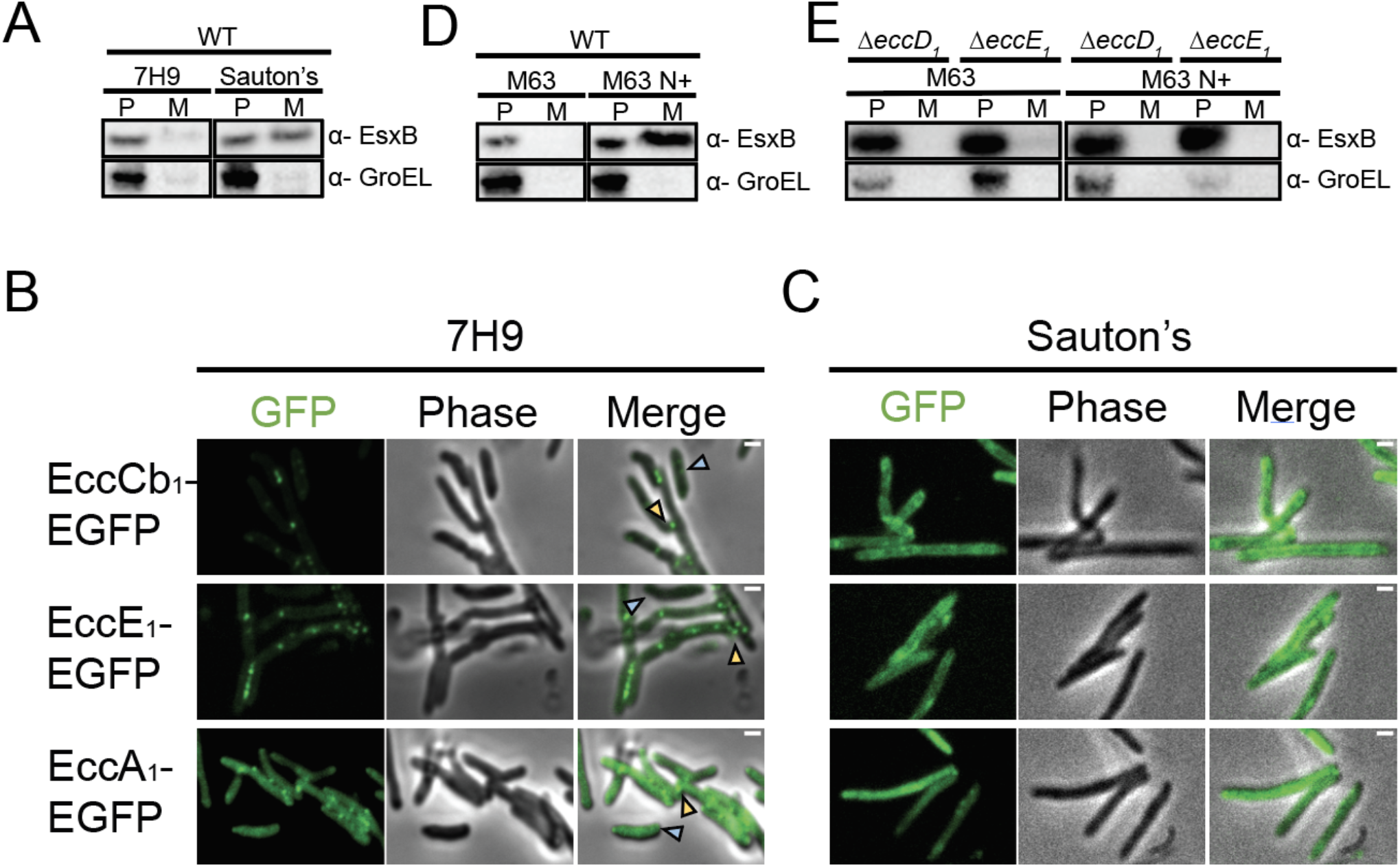
EsxB secretion is triggered by environmental nitrogen levels. Western blot shows the ESX-1 substrate EsxB secreted into the medium. P represents pellets and M represents medium, and anti-GroEL antibody shows the integrity of the loaded samples. A) Secretion assay in 7H9 and Sauton’s medium. Experiments shown are representative of three biological replicates. B) EccCb_1_-EGFP, EccE_1_-EGFP, and EccA_1_-EGFP cultured in 7H9 medium are shown. Yellow arrows depict foci at cell-cell contacts, blue arrows depict singular cells lacking foci. C) EccCb_1_-EGFP, EccE_1_-EGFP, and EccA_1_-EGFP cultured in Sauton’s medium are shown. Scale bar located on top right corner of images, 1µm. D) Secretion assay in defined minimal medium, M63 and M63 with ammonium chloride (M63 N+). E) Secretion assay with M63 and M63 N+ on ESX-1 *eccD*_*1*_ and *eccE*_*1*_ knockouts. Secretion requires the entire ESX-1 assembly to occur.

The two media differ in the amount of available nitrogen and carbon sources, with Sauton’s medium containing about 6X as much nitrogen and 25X as much carbon as 7H9 medium. (7H9 contains 1.1 mM elemental nitrogen and 10.4 mM elemental carbon, while Sauton’s contains 6.4 mM and 269.8 mM, respectively) Higher concentrations of carbon favor typical, clumped growth of *M. smegmatis* in liquid medium (DePas et al., 2019), while higher concentrations of nitrogen favor planktonic growth (DePas et al., 2019; Glaeser and Taylor, 1978). Thus, we investigated whether altering the concentration of nitrogen alone is sufficient to induce ESX-1 secretion of EsxB into the medium. Cultures were grown in modified M63 minimal medium, which supports mycobacterial growth in a range of carbon and/or nitrogen concentrations (DePas et al., 2019). We systematically altered the level of nitrogen from 1.02 mM nitrogen (M63) to 6.26 mM (M63 N+) characteristic respectively of 7H9 and Sauton’s. Secretion of the ESX-1 substrate, EsxB, into the spent media only occurred in the M63 N+ medium (Figure 3B). We confirmed that this secretion of EsxB was dependent on the ESX-1 system (Figure 3C). We next tested whether cells lacking ESX-1 show clumping in low nitrogen (M63) and dispersal in high nitrogen (M63 N+). Cultures were analyzed by macroscopic analysis of growths in a culture tube, as reported prior(DePas et al., 2019). The growths showed that deletion mutants of ESX-1 did not have differential growth from wild-type cells in either medium (Supplemental Figure 4).

Given the strong induction of ESX-1 secretion we asked whether the transition of ESX-1 to a secretion-competent form is due to changes in gene expression of ESX-1 components. Our findings show that expression levels of EccCb_1_-EGFP are similar in M63 and M63 N+ (Figure 4A). To investigate transcriptional changes in ESX-1 we analyzed global transcriptional profiles (RNAseq) of *M. smegmatis* grown in M63 and M63 N+ media (Figure 4B). Analysis of the genes in the ESX-1 operon showed these (MSMEG_0055 – MSMEG_0082) are not differentially translated (Figure 4B, orange circles). In addition, we observed consistent signatures for the major downregulated genes in high nitrogen conditions, such as nitrogen importers (Figure 4B, green circles; MSMEG_2425-AmtB, MSMEG_6259-Amt1, MSMEG_4635-AmtA). Collectively, these results strongly suggest that high nitrogen promotes ESX-1 secretion. through post-translational changes to the system

**Figure 4:**
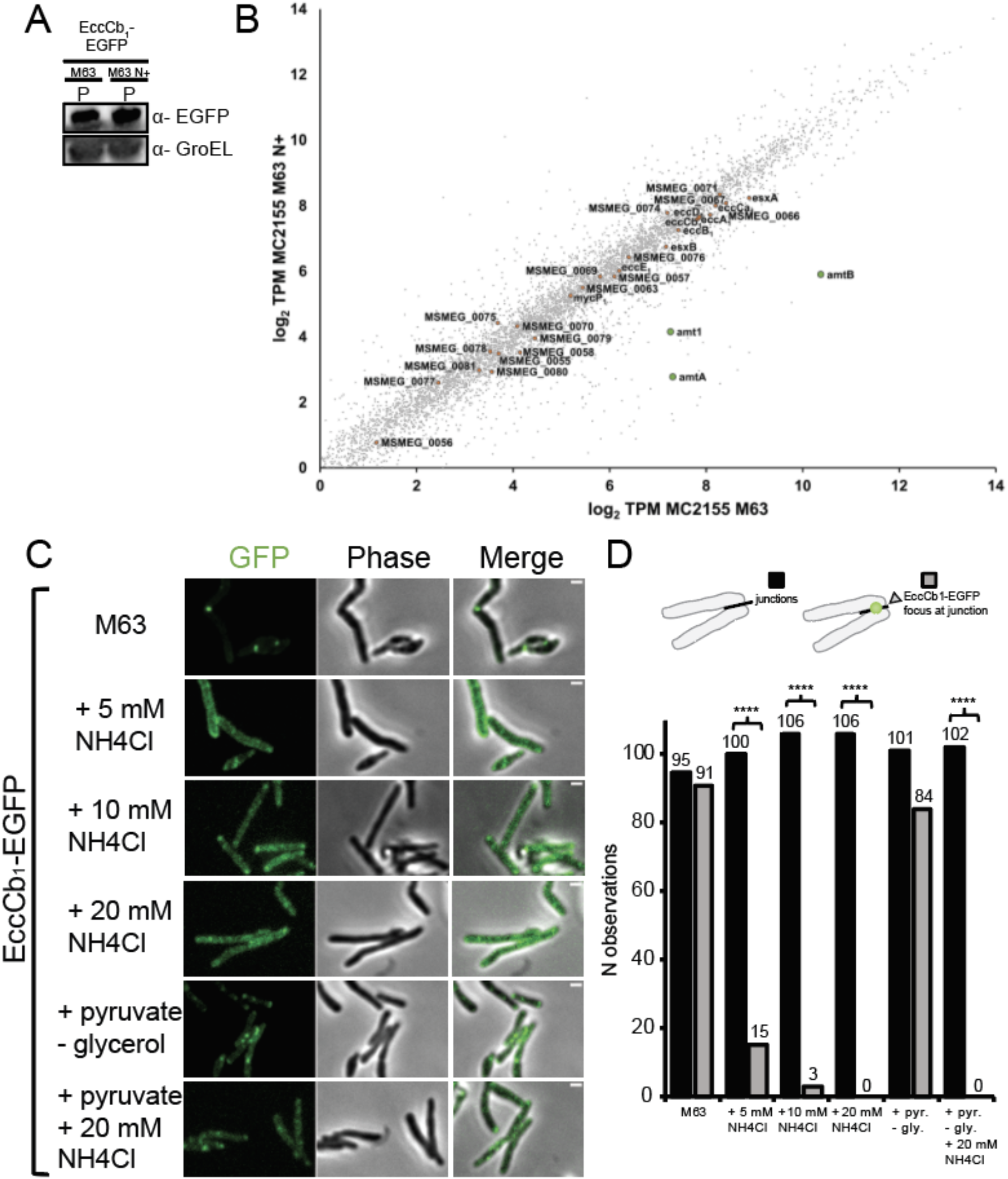
ESX-1 foci are inhibited by high nitrogen in the medium. A) ESX-1 expression in cells was assessed with an endogenous EGFP marker on EccCb_1_. Anti-GFP demonstrates levels of ESX-1 expression in M63 and M63 N+ medium. B) Transcript abundances (as log_2_ transcripts per million) measured by RNAseq in wild-type *M. smegmatis* growing in M63 (y-axis) vs M63 N+ (x-axis). Orange circles represent ESX-1 genes. Green circles represent downregulated transporters in response to nitrogen addition. Time averaged confocal images acquired every 5 minutes for the duration of an hour are depicted. GFP, Phase and merged channels are shown. Scale bar located on top right corner of images, 1µm. A) EccCb_1_-EGFP, EccE_1_-EGFP, and EccA_1_-EGFP cultured in 7H9 medium are shown. Yellow arrows depict foci at cell-cell contacts, blue arrows depict singular cells lacking foci. B) EccCb_1_-EGFP, EccE_1_-EGFP, and EccA_1_-EGFP cultured in Sauton’s medium are shown. C) EccCb_1_-EGFP cultured in M63, M63 N+, or M63 minimal medium supplemented with pyruvate to induce carbon scarcity. D) Foci were quantified across 90+ contacts per strain for three biological triplicates in distinct M63 media. Exact numbers are denoted above corresponding data bar and are outlined on the Y-axis as N observations. Difference between both measurements is statistically significant per student’s t-test, **** P < 0.0001.

### Nitrogen levels regulate ESX-1 foci formation

To investigate if nitrogen was sufficient to trigger the dissolution of ESX-1 foci, we imaged EccCb_1_-EGFP grown in M63 supplemented with varying concentrations of nitrogen. EccCb_1_-EGFP cultures grown in M63 medium exhibited foci formed at 100% of cell-cell contacts, similar to those observed in 7H9 medium (Figure 4C and 4D). Stepwise addition of NH_4_Cl to the growth medium led to dissipation of foci as a function of NH_4_Cl concentration; at 5 mM NH_4_Cl 15% of contacts retained focus formation, at 10 mM NH_4_Cl 3% of contacts retained focus formation and at 20 mM NH_4_Cl 0% of contacts exhibited focus formation (Figure 4C and 4D).

As high nitrogen conditions induce planktonic growth (DePas et al., 2019), we investigated if other stimuli which trigger planktonic growth of cells, such as carbon starvation, also regulate focus formation. We cultured cells in M63 medium in which pyruvate was provided as a less bioavailable carbon source compared to glycerol (M63 + pyruvate / - glycerol)(DePas et al., 2019). Time averaged images of pyruvate grown cultures demonstrated that focus formation occurred at 81% of cell-cell contacts (Figure 4C and D). In media with both pyruvate and nitrogen (M63 + pyruvate - glycerol + 20 mM NH_4_Cl), none of the cells exhibited stable foci at cell-cell contacts (Figure 4C and 4D). We note that in pyruvate supplemented culture conditions, we sometimes observed ESX-1 at poles in isolated cells. It is possible these cells are experiencing additional stress causing this localization. Overall, our experiments demonstrate that ESX-1 responds specifically to excess nitrogen by dissociating ESX-1 foci at cell-cell contacts. This change in ESX-1 may be in response to nitrogen levels rather than shift to planktonic growth. Further, the change in localization correlated with active secretion of EsxB into the spent growth medium. This distinct change indicated that when actively secreting, ESX-1 is dispersed from its focal form, which strongly suggested that formation of ESX-1 foci correlates with an alternate non-secretory state of ESX-1.

### Transcriptional response to nitrogen reveals a function of ESX-1 in SOS regulation

To probe the possible function of ESX-1 in nitrogen response, we analyzed RNAseq profiles of various strains of *M. smegmatis* grown in either M63 or M63 N+. Using a series of *M. smegmatis* strains (wild-type, Δ*esxB*, Δ*eccCa*_*1*_, and Δ*eccD*_*1*_) we assessed whether nitrogen-dependent transcriptional gene regulation of genes by nitrogen is mediated by ESX-1. First, we examined a set of genes previously described in nitrogen metabolism (Amon et al., 2009; Petridis et al., 2015; Williams et al., 2013). Consistent with previous studies, the transcriptional profiles exhibited downregulation of nitrogen importer genes (*amtB, amt1* and *amtB*), the GlnR operon and nitrate assimilation genes, in M63 N+ (Figure 5A). The differential transcription of these genes remained unchanged in our ESX-1 deletion strains, which was confirmed by RT-qPCR (Figure 5B). Thus, ESX-1 was not required for the general transcriptional response to nitrogen (Figure 5A).

**Figure 5:**
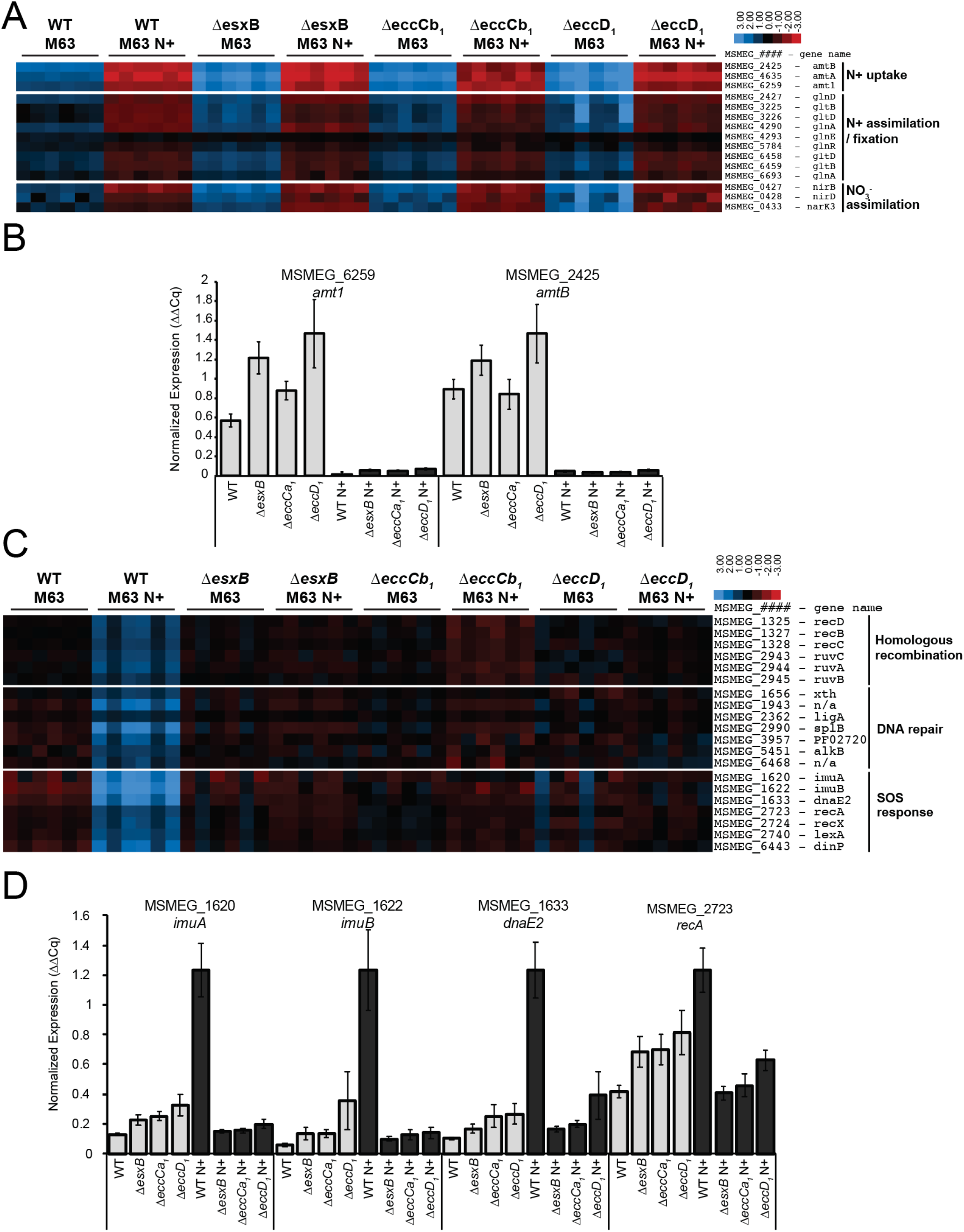
Transcriptional responses to nitrogen reveal a function of ESX-1 in SOS response. A) Heatmap demonstrating ESX-1 knockouts do not influence cellular response to nitrogen. Log-2 scale for the heatmap is shown in the upper right hand corner of the heatmap. Data are representative of 3 biological replicates and 2 technical replicates. B) RT-qPCR of *amt1* (left) and *amtB* (right) transporters in distinct strains. Results were normalized to sigA and reported as ΔΔCq. Normalized gene expression across distinct *M. smegmatis* strains grown in two M63 media. Wildtype (WT), ΔesxB, and ΔeccD_1_ are shown. N = 3 experiments. C) Heatmap rendered from RNAseq data displaying DNA repair and SOS response elements upregulated in response to M63 N+. Log-2 scale for the heatmap is shown in the upper right hand corner of the heatmap. Data are representative of 3 biological replicates and 2 technical replicates. D) RT-qPCR of SOS elements: *imuA, imuB, dnaE2, recA*. Results were normalized relative to *sigA*. Data are represented as ΔΔCq. Normalized gene expression across distinct *M. smegmatis* strains grown in two M63 media. Wildtype (WT), ΔesxB, and ΔeccD_1_ are shown. N = 3 experiments.

Further analysis of RNAseq data from *M. smegmatis* strains grown in M63 and M63 +N showed an unexpected effect of ESX-1 on pathways not previously linked to nitrogen metabolism and ESX-1 functions. Overall, growing wild-type cultures in excess nitrogen resulted in 342 downregulated and 438 upregulated genes in response to excess nitrogen (plotted in Supplemental Figure 5). Genes were considered differentially expressed if they had an absolute log_2_ fold change greater than 1 and were significantly differential at a false discovery rate (FDR) of 5%. In contrast, the ESX-1 mutants showed a distinct profile with 353 downregulated genes and 526 upregulated genes in Δ*eccCa*_*1*_, 389 downregulated and 621 upregulated genes in Δ*eccD*_*1*_, and 351 downregulated genes and 607 upregulated genes in Δ*esxB*, (plotted in Supplemental Figure 5). The differences in the transcriptional profile of the mutants were clustered in known regulons. Most strikingly, cultures grown in M63 N+ exhibited a strong upregulation of error-prone DNA replication pathways, such as the LexA regulon and associated genes in the SOS response; this upregulation was absent in ESX-1 mutants affecting membrane complex formation (Δ*eccCa*_*1*_ and Δ*eccD1*) and in the ESX-1 secreted substrate (Δ*esxB*) (Figure 5C). These observations were validated using RT-qPCR on error prone DNA replication machinery genes, *imuA, imuB, dnaE2*, and *recA* (Figure 5D). Interestingly, baseline levels of these elements were generally higher in ESX-1 knockout cultures grown in M63 medium, which is especially evident in *recA*. This suggests ESX-1 mediates the adaptational response to environmental triggers, such as nitrogen. In the absence of ESX-1, a growing culture might not sense environmental conditions fully and compensates by harboring a higher baseline level of SOS induced error prone DNA replication machinery.

## Discussion

Environmental bacteria live in nutritionally dynamic conditions and require mechanisms to sense and respond to local overgrowth. Our studies show that ESX-1 alters its function in response to nitrogen levels. At low nitrogen levels, cells do not secrete; instead ESX-1 assembles into a megacomplex across spanning both plasma membranes within minutes of cell-cell contact (Figure 1). At high nitrogen levels, cells secrete ESX substrates, and the megacomplexes disap-pears. ESX-1 does not appear to be required for forming cell-cell contacts themselves, as ESX-1 mutants still clump (Supplemental Figure 4). Rather, ESX-1 may contribute to sensing cell contacts to regulate downstream signaling pathways.

The presence of ammonium chloride in the medium can be seen as a model of increased nitrogenous waste from increased bacterial growth and density (DePas et al., 2019; Vince et al., 1973). Based on our results, we propose a model for ESX-1 secretion throughout various growth phases (Figure 6). When the bacteria are in initial growth phases in nutritionally favorable conditions, the nitrogenous waste levels are low, and the bacteria grow in aggregates. This aggregation triggers accumulation of ESX-1 systems at the cell-to-cell contacts, which in turn activates ESX-1 focus formation. As a culture enters stationary phase, nitrogenous waste accumulates and the bacterial surface changes to promote dissociation of the cells, loss of cell-to-cell contact and diffusion of ESX-1 systems in the membrane. These changes trigger ESX-1 secretion. Although the downstream consequences of ESX-1 secretion are likely pleotropic, one clear outcome is the upregulation of genes implicated in SOS response required for translesion synthesis and mutagenic replication (Figure 5). We speculate that this upregulation leads to an increase in mutagenic replication and potentially generation of multiple phenotypes to disperse into new environments during planktonic growth (Kivisaar, 2003).

**Figure 6:**
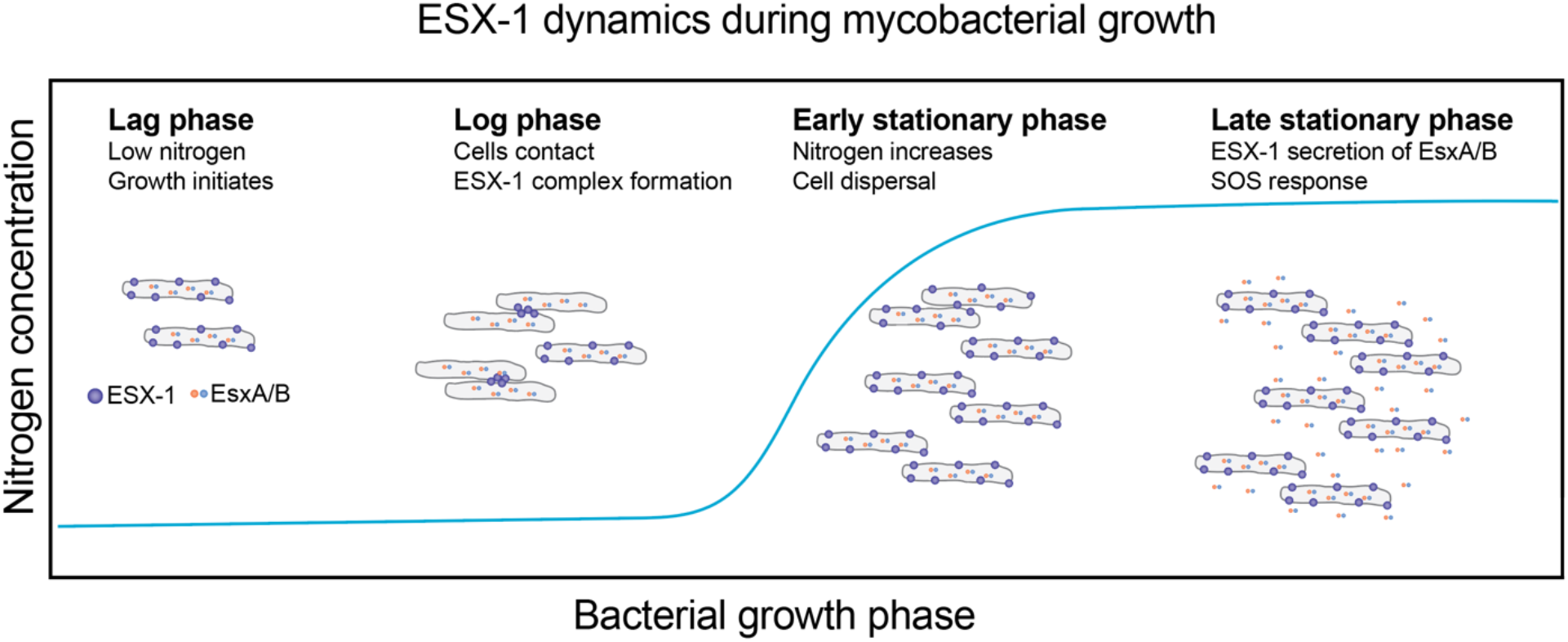
Schematic of ESX-1 dynamics during mycobacterial growth. As a nascent culture grows, ESX-1 is dispersed throughout the plasma membrane of planktonic cells. As growth continues, cells clump and ESX-1 forms complexes at cell-cell contact points. Once nitrogen levels start increasing in the culture cells commence dispersion and ESX-1 complexes dissipate from cell-cell contact points. Once the nitrogen levels are saturated, ESX-1 secretion of substrates EsxA/B commences and the SOS response is upregulated in cells.

*M. smegmatis* ESX-1 has been most extensively studied in the context of direct conjugal transfer, where it is a key negative regulator of the process in a donor strain (Cao et al., 2015; Coros et al., 2008; Derbyshire and Gray, 2014; Gray et al., 2016). How our findings relate to conjugation is not yet clear, as conjugation studies will require optimization of multiple strains beyond the scope of this work. The formation of ESX-1 foci at cell-cell contacts is certainly consistent with a role in conjugation at junctions but we speculate that ESX-1 may contribute to cell-cell communication in a broader sense ESX-1 is not required for DNA transfer per se, and it has a negative inhibitory role in selecting a specific acceptor strain. Its role in SOS response may also be seen as part of conjugation preparations for horizontal gene transfer, as the donor cell is preparing its genome for rapid transfer of DNA to an acceptor strain (Guerin et al., 2009). In this light, DNA replication induced by the error prone polymerase, DnaE2, would trigger production of DNA to prepare the donor strain for DNA transfer. In the absence of ESX-1, conjugation is less regulated in the donor strain; any given donor strain is prepared for rapid DNA transfer to an acceptor strain (Derbyshire and Gray, 2014; Gray et al., 2016). This may be caused in part by the constitutive increase in basal transcription of SOS response elements (Figure 5), which as such could represent preparation for conjugation (Flint et al., 2004).

Our findings have implications for the roles of ESX-1 in virulence of *M. tuberculosis*. It has generally been assumed that ESX-1 secretes virulence factors that promote growth in the host intracellular environment (Gröschel et al., 2016). However, the concept of the secretion of a direct toxin or effector is hard to reconcile with the mild phenotypic differences seen at the early stages of experimental infection. It has been hypothesized that ESX-1 secretion may only be important for virulence in the chronic stages of infection (Stanley et al., 2003). Our work suggests one role of ESX-1 may be to regulate the mycobacterial response to the accumulation of nitrogenous waste in the phagosome (Gordon et al., 1980), a condition that is also known to cause phagosomal arrest. We speculate ESX-1 mutants may be more rapidly cleared from macrophages because they cannot adapt to the nutrient starved, bacterial-dense environment of the phagosome. A role for this type of adaptive evolution was postulated in seminal work documenting the importance of translesion synthesis in long term survival and adaptation in *M. tuberculosis* (Boshoff et al., 2004). That work showed that mutations induced by DnaE2 are required for the long-term survival of mycobacteria in a murine model and hypothesized that adaptive evolution is responsible for this effect. Further study will be needed to show that the virulence phenotype of *M. tuberculosis* ESX-1 is related to these findings in a mycobacterial model organism.

## Materials and Methods

### Bacterial culture and growth conditions

*M. smegmatis* cultures were grown in Difco 7H9 medium or modified M63 medium (DePas et al., 2019), at 37 °C and 150 RPM. Starter cultures were inoculated from a single colony on a 7H9 plate and grown for 72 hours, and then diluted 1:50 to inoculate subsequent cultures. Where appropriate, hygromycin was used at a final concentration of 100 µg/mL and kanamycin was used at a final concentration of 25 µg/mL. To make M63 N+, NH_4_Cl was added to the medium to a final concentration of 20 mM, unless otherwise noted. All cultures contained 0.05% tween, unless otherwise noted.

*E. coli* cultures were grown in LB medium for plasmid amplification and TB medium for protein overexpression. Cultures were incubated at 37 °C, 150 RPM. Where appropriate, kanamycin was used at a final concentration of 50 µg/mL, and hygromycin at a final concentration of 150 µg/mL. Protein expression was induced at an OD_600nm_ of 1.0-1.2 by addition of 1 µM IPTG for 2 hours.

### Generation of mutant strains and complementing constructs

To construct chromosomally labeled EGFP strains we used the ORBIT method (Murphy et al., 2018). In brief, pKM444 was introduced into wild-type and ESX-1 knockouts of MC^2^155 *M. smegmatis*, resulting in a strain producing annealase and resolvase under a P_tet_ promoter. Targeting plasmid pKM468 and gene targeting ultramers (Supplemental Table 1) were electroporated. Recovered cells were plated on 7H9 hygromycin plates. Colonies were screened for hygromycin resistance and verified using colony PCR and sanger sequencing. Complementation studies were made by introducing the complementing gene of interest upstream of the mop promoter in plasmid pMV306. pMV306 was restriction digested at the DraI site and inserts were amplified by PCR with 20bp overlaps with cut pMV306. Infusion was used to assemble the plasmid. Plasmids were electroporated into cells containing the corresponding deletions for complementation studies.

For co-culture studies, we introduced a cytoplasmic mCherry under a groEL promoter in pMV261 into either wild-type and ESX-1 knockouts of MC^2^155. pMV261 was amplified by PCR, upstream of the groEL2 promoter, mCherry was amplified by PCR with 20bp overlaps to pMV261 and the final product was assembled by infusion. All constructed strains, primers and plasmids are reported in Supplementary Table 1.

### Live cell confocal microscopy

*M. smegmatis* cultures were grown in liquid cultures to exponential phase (OD_600 nm_ between 0.6-0.8) and placed into microchannels (ibidi µ-slide VI 0.4 slides: Ibidi 80606, Ibiditreat #1.5) which had been coated with PDMS prior to imaging. PDMS was prepared by mixing a 1:10 ratio of curing agent to PDMS and applied to the channels. Excess PDMS was removed from the channels by blowing air through individual channels, then the PDMS was cured by incubation in an oven set to 80°C for 20 minutes. Cells were allowed to settle to the bottom of the channel for 10 minutes and then washed with appropriate medium. The chamber was kept at 37°C in a temperature-controlled enclosure (OkoLab) throughout imaging.

Cells were imaged on a spinning disc confocal system with a 488 nm excitation laser at 50 ms exposure at 0.1 Hz (30% power), with a 100x objective (Nikon Ph3 100x N.A. 1.4) on a Nikon TI-E stand equipped with a spinning-disk confocal head (CUS10, Yokogawa) and an EM-CCD camera (Hammamatsu C9100-13). Images were generally captured in 5-minute intervals for 1 hour. This interval was determined to be the most appropriate when taking into consideration the 4-hour doubling time for *M. smegmatis*, to fully capture the dynamics of the foci. All imaging was done across biological triplicates. Images were analyzed by manually outlining the cell-cell junctions and measuring EGFP intensity for the contact site. During analysis, data were single blinded.

### Secretion assays

Cultures were inoculated into the appropriate medium from a colony growing on a 7H9 plate. The cultures were then transferred and grown to mid-log phase (OD_600 nm_ ∼0.6-0.8). A second transfer of the cultures was done into medium lacking tween and allowed to reach an OD_600 nm_ of 0.8. Cells were harvested by centrifugation at 3000 RPM in an Eppendorf centrifuge. The supernatant was filtered through a 0.22 µm filter and concentrated 500-fold using a 3,000 MWCO Amicon ultra-15 filter (25 mL to 50 µL). Whole cell pellets were re-suspended in PBS. Whole cell resuspensions and culture concentrates were incubated with SDS loading dye and analyzed by SDS-PAGE using Invitrogen Bis Tris SDS gels. For western blotting of *M. smegmatis* EsxB a polyclonal mouse antibody was raised using an EsxB antigenic peptide sequence. The anti-GroEL antibody produced in rabbit (Sigma G6532-.5ML) was used to represent sample integrity.

### RNAseq sample preparation

RNA was extracted from cultures grown to OD_600 nm_ ∼ 0.7-0.8. Three biological replicates (single colonies) were prepared for every strain assayed. The NEB RNA extraction kit was used as directed except for the lysis step, which was completed by bead beating the cells with 0.1 mm zirconia beads. To deplete rRNA, we used the NEB rRNA kit as directed. Samples were prepared using the NEBNext Ultra RNA library prep kit for Illumina, and barcoding oligos were used to pool the libraries. Two technical replicates were prepared for analysis. The resulting pools were sequenced with the Illumina NovaSeq in 75 nt single end reads, resulting in 15M reads per sample. All constructed strains are reported in Supplementary Table 1.

### RT-qPCR sample preparation

Primers used for RT-qPCR were validated by qPCR using serial dilutions of genomic DNA extracted from *M. smegmatis* MC^2^155. The NEB Luna® qPCR Master Mix was used as directed by the manufacturer. For RT-qPCR, RNA was prepared as outlined in the RNAseq section, in biological triplicates. Samples were normalized to 75 ng/µL and assayed in technical duplicates analyzed using the NEB Luna ® Universal One-Step RT-qPCR kit. All constructed strains and primers are reported in Supplementary Table 1.

### Bioinformatics

Transcripts were pseudo aligned with KALLISTO (Bray et al., 2016) to the *M. smegmatis* MC^2^155 coding sequences (NCBI accession GCF_000015005.1) to estimate relative abundances (reported as TPM values) and estimated counts (est_counts). Further analysis was restricted to transcripts with TPM ≥ 1 in at least one sample. Differential expression between different genotypes and growth conditions was estimated using LIMMA version 3 (Ritchie et al., 2015; Smyth, 2004), and transcripts were considered to be significantly differentially expressed if they had a log2 fold change of at least 1 at a false discovery rate (FDR) of 5%. Corresponding files have been deposited in the Gene Expression Omnibus database under accession number GSE185010.

## Acknowledgements

We thank Jeff Cox for providing strains of *M. smegmatis* MC^2^155 Δ*eccB*_*1*_, Δ*eccCb*_*1*_, Δ*eccD*_*1*_, and Δ*eccE*_*1*_. We are grateful to the Infectious Disease Team (Amy Lyden, Emily Crawford) and the Sequencing Team (Norma Neff, Feiqiao Brian Yu, Michelle Tan, Angela Detweiler, Honey Mekonen) at the Chan Zuckerberg Biohub who assisted with RNAseq sample preparation guidelines and sequencing. We thank Carol A. Gross for helpful discussions and critical reading of the manuscript. We acknowledge the Burroughs Wellcome Fund Postdoctoral Enrichment Award #1019894, NIH T32 training grant (5T32HL007185-42) and the University of California President’s Postdoctoral Fellowship to NH; and NIH NIAID R01AI128214 and Chan Zuckerberg Biohub funding to OSR.

## Contributions

NH and OSR designed the project and wrote the initial manuscript. NH and RN designed and engineered the *M. smegmatis* strains made for this study. NH, PDO, and FC designed and implemented the imaging experiments. NH, MV, and AS designed and implemented the RNAseq data acquisition and analysis. All authors revised and edited the manuscript.

---

**Supplemental Figure 1:**
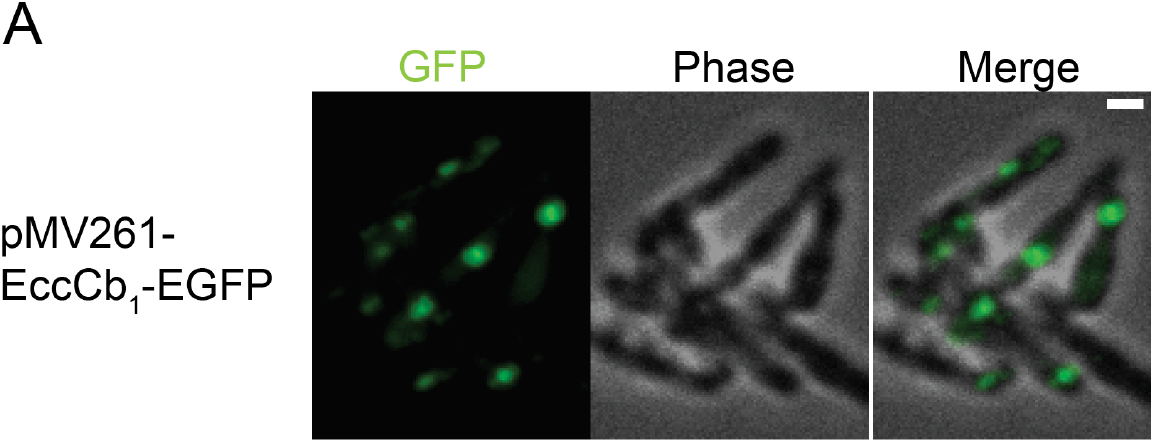
Visualization of EGFP constructs expressed on a plasmid. A) Time averaged confocal images acquired every 5 minutes for the duration of an hour are depicted. Phase and GFP channels are merged. Scale bar located on top right corner of image, 1 µm. Images were captured on cells expressing EccCb_1_-EGFP on 7H9 medium. Images representative of three biological replicates.

**Supplemental Figure 2:**
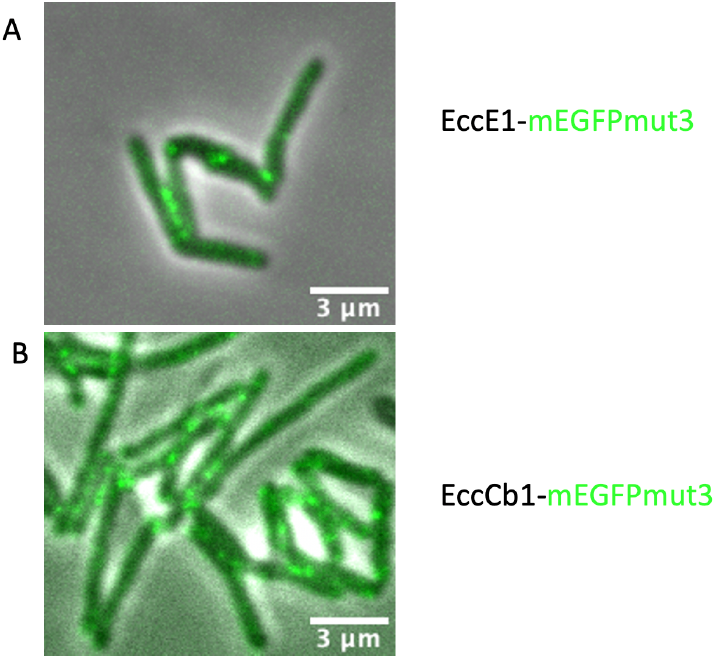
Visualization of mEGFPmut3 constructs. Time averaged confocal images acquired every 5 minutes for the duration of an hour are depicted. Phase and GFP channels are merged. Scale bar located on top right corner of images, 3 µm. Strains were imaged in 7H9 medium A) EccCb_1_ – mEGFP mut3 B) EccE_1_ – mEGFP mut3 is shown. Images representative of three biological replicates.

**Supplemental Figure 3:**
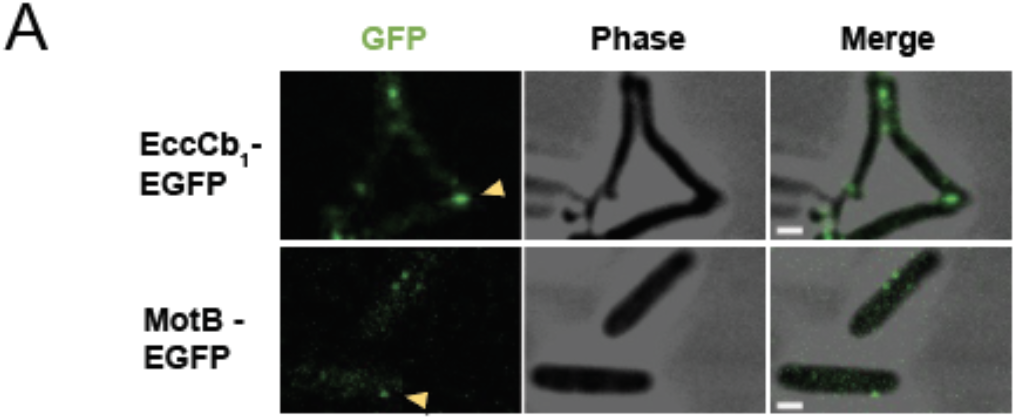
Images of single acquisitions of EGFP tagged ESX-1 and MotB. A) Representative confocal microscopy images of single acquisitions in *M. smegmatis* EccCb_1_-EGFP (top) and *E. coli* MotB-EGFP. Foci are delineated by yellow arrowheads. Images representative of three biological replicates.

**Supplemental Figure 4:**
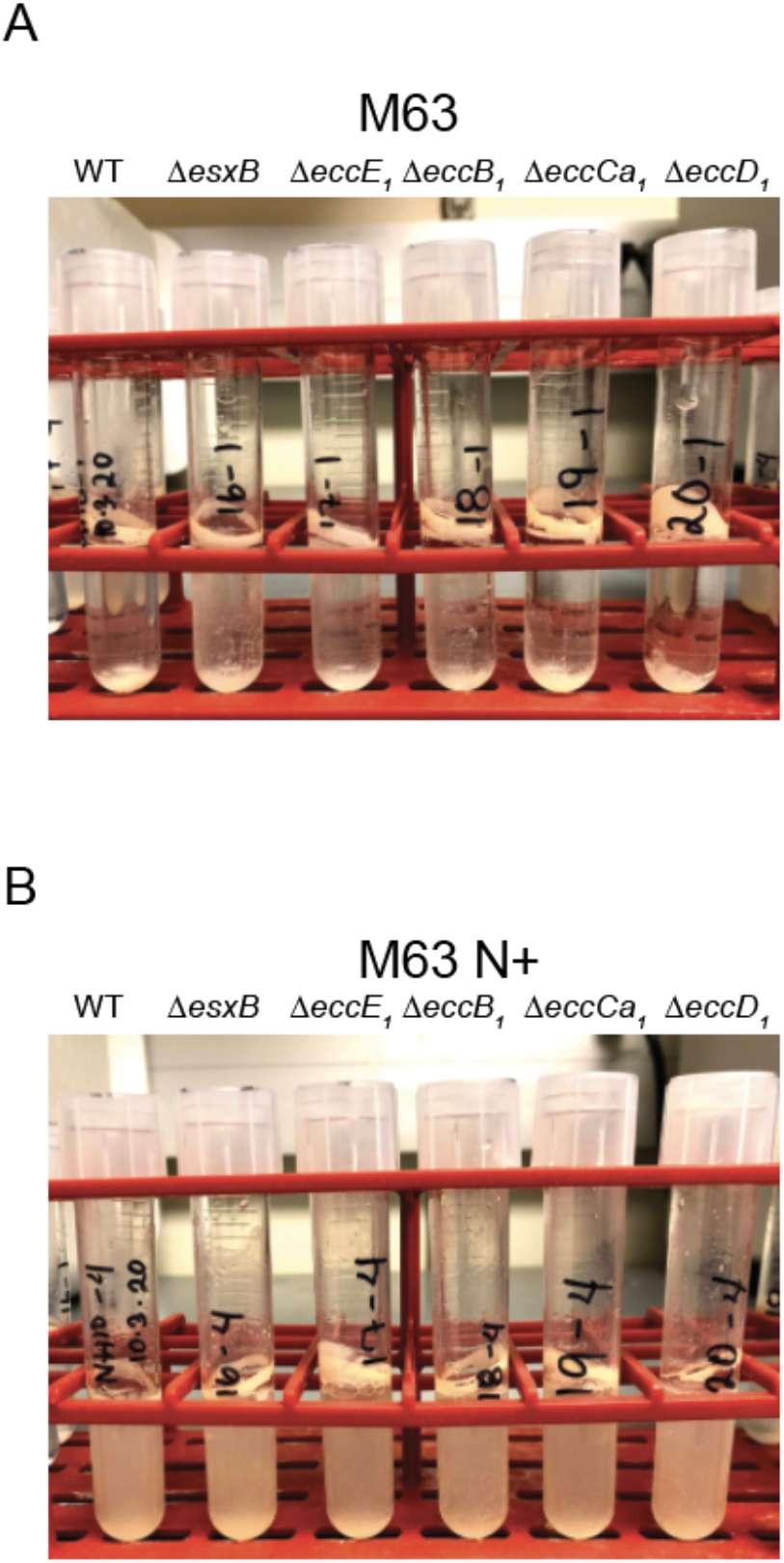
*M. smegmatis* cultures in M63 vs M63 N+. Wildtype (WT), ΔesxB, ΔeccE_1_, ΔeccB_1_, ΔeccCa_1_, and ΔeccD_1_ are shown. Representative of three biological replicate growths. A) Growth in M63 medium. B) Growth in M63 N+ medium.

**Supplemental Figure 5:**
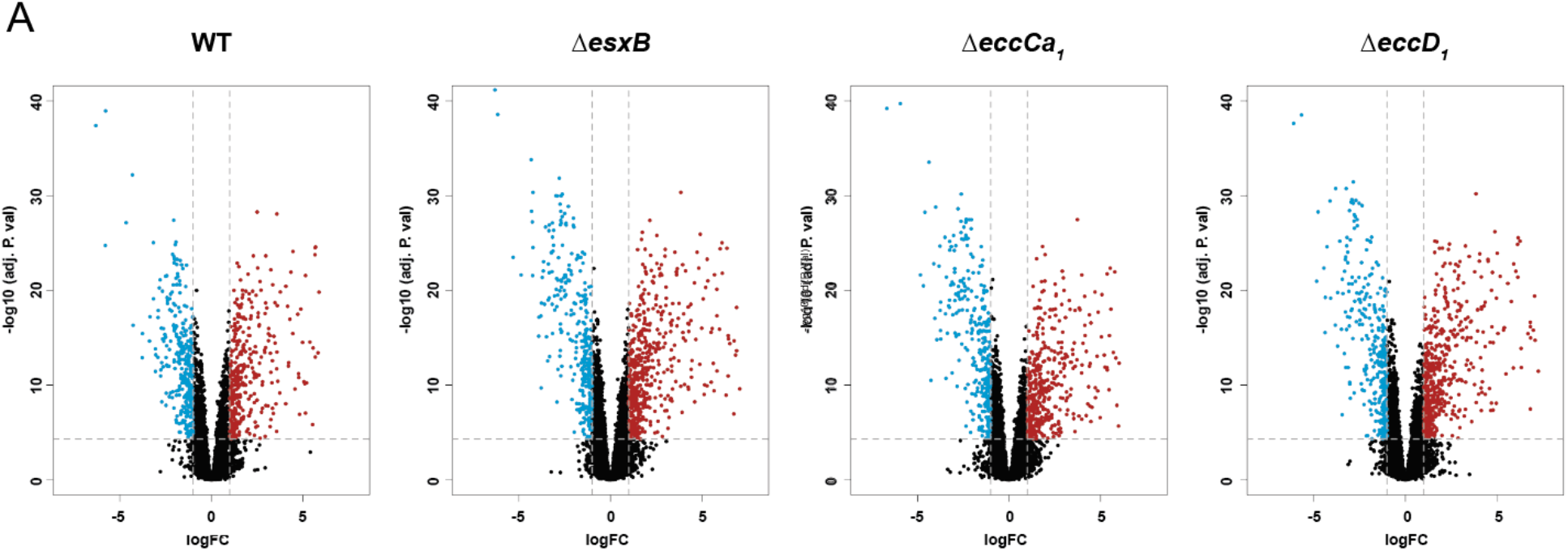
Differential expression in distinct *M. smegmatis* strains in M63 vs M63 N+. A) Normalized gene expression across distinct *M. smegmatis* strains grown in two M63 media. Wildtype (WT), ΔesxB, and ΔeccD_1_ are shown. Data are representative of 3 biological replicates and 2 technical replicates. A) Differential expression volcano plots of *M. smegmatis* strains, WT, followed by ESX-1 knockouts. Blue genes are upregulated, red genes are downregulated.

## Notes

### Competing Interest Statement

The authors have declared no competing interest.

## References

Amon, J., Titgemeyer, F., Burkovski, A., 2009. A genomic view on nitrogen metabolism and nitrogen control in mycobacteria. J Mol Microbiol Biotechnol 17, 20–29. doi:10.1159/000159195

Baptista, C., Barreto, H.C., São-José, C., 2013. High levels of DegU-P activate an Esat-6-like secretion system in Bacillus subtilis. PLoS One 8, e67840. doi:10.1371/journal.pone.0067840

Beckham, K.S.H., Ciccarelli, L., Bunduc, C.M., Mertens, H.D.T., Ummels, R., Lugmayr, W., Mayr, J., Rettel, M., Savitski, M.M., Svergun, D.I., Bitter, W., Wilmanns, M., Marlovits, T.C., Parret, A.H.A., Houben, E.N.G., 2017. Structure of the mycobacterial ESX-5 type VII secretion system membrane complex by single-particle analysis. Nat. Microbiol. 2, 17047. doi:10.1038/nmicrobiol.2017.47

Beckham, K.S.H., Ritter, C., Chojnowski, G., Ziemianowicz, D.S., Mullapudi, E., Rettel, M., Savitski, M.M., Mortensen, S.A., Kosinski, J., Wilmanns, M., 2021. Structure of the mycobacterial ESX-5 type VII secretion system pore complex. Sci. Adv. 7. doi:10.1126/sciadv.abg9923

Berthet, F.X., Rasmussen, P.B., Rosenkrands, I., Andersen, P., Gicquel, B., 1998. A Mycobacterium tuberculosis operon encoding ESAT-6 and a novel low-molecular-mass culture filtrate protein (CFP-10). Microbiology (Reading, Engl.) 144 (Pt 11), 3195–3203. doi:10.1099/00221287-144-11-3195

Boshoff, H.I.M., Myers, T.G., Copp, B.R., McNeil, M.R., Wilson, M.A., Barry, C.E., 2004. The transcriptional responses of Mycobacterium tuberculosis to inhibitors of metabolism: novel insights into drug mechanisms of action. J. Biol. Chem. 279, 40174–40184. doi:10.1074/jbc.M406796200

Bray, N.L., Pimentel, H., Melsted, P., Pachter, L., 2016. Near-optimal probabilistic RNA-seq quantification. Nat. Biotechnol. 34, 525–527. doi:10.1038/nbt.3519

Burts, M.L., DeDent, A.C., Missiakas, D.M., 2008. EsaC substrate for the ESAT-6 secretion pathway and its role in persistent infections of Staphylococcus aureus. Mol. Microbiol. 69, 736–746. doi:10.1111/j.1365-2958.2008.06324.x

Burts, M.L., Williams, W.A., DeBord, K., Missiakas, D.M., 2005. EsxA and EsxB are secreted by an ESAT-6-like system that is required for the pathogenesis of Staphylococcus aureus infections. Proc. Natl. Acad. Sci. USA 102, 1169–1174. doi:10.1073/pnas.0405620102

Cao, G., Howard, S.T., Zhang, P., Wang, X., Chen, X.-L., Samten, B., Pang, X., 2015. EspR, a regulator of the ESX-1 secretion system in Mycobacterium tuberculosis, is directly regulated by the two-component systems MprAB and PhoPR. Microbiology (Reading, Engl.) 161, 477–489. doi:10.1099/mic.0.000023

Carlsson, F., Joshi, S.A., Rangell, L., Brown, E.J., 2009. Polar localization of virulence-related Esx-1 secretion in mycobacteria. PLoS Pathog. 5, e1000285. doi:10.1371/journal.ppat.1000285

Coffman, V.C., Wu, J.-Q., 2012. Counting protein molecules using quantitative fluorescence microscopy. Trends Biochem. Sci. 37, 499–506. doi:10.1016/j.tibs.2012.08.002

Cole, S.T., Brosch, R., Parkhill, J., Garnier, T., Churcher, C., Harris, D., Gordon, S.V., Eiglmeier, K., Gas, S., Barry, C.E., Tekaia, F., Badcock, K., Basham, D., Brown, D., Chillingworth, T., Connor, R., Davies, R., Devlin, K., Feltwell, T., Gentles, S., Hamlin, N., Holroyd, S., Hornsby, T., Jagels, K., Krogh, A., McLean, J., Moule, S., Murphy, L., Oliver, K., Osborne, J., Quail, M.A., Rajandream, M.A., Rogers, J., Rutter, S., Seeger, K., Skelton, J., Squares, R., Squares, S., Sulston, J.E., Taylor, K., Whitehead, S., Barrell, B.G., 1998. Deciphering the biology of Mycobacterium tuberculosis from the complete genome sequence. Nature 393, 537–544. doi:10.1038/31159

Converse, S.E., Cox, J.S., 2005. A protein secretion pathway critical for Mycobacterium tuberculosis virulence is conserved and functional in Mycobacterium smegmatis. J. Bacteriol. 187, 1238–1245. doi:10.1128/JB.187.4.1238-1245.2005

Coros, A., Callahan, B., Battaglioli, E., Derbyshire, K.M., 2008. The specialized secretory apparatus ESX-1 is essential for DNA transfer in Mycobacterium smegmatis. Mol. Microbiol. 69, 794–808. doi:10.1111/j.1365-2958.2008.06299.x

DePas, W.H., Bergkessel, M., Newman, D.K., 2019. Aggregation of Nontuberculous Mycobacteria Is Regulated by Carbon-Nitrogen Balance. MBio 10. doi:10.1128/mBio.01715-19

Derbyshire, K.M., Gray, T.A., 2014. Distributive Conjugal Transfer: New Insights into Horizontal Gene Transfer and Genetic Exchange in Mycobacteria. Microbiol. Spectr. 2. doi:10.1128/microbiolspec.MGM2-0022-2013

Dumas, E., Christina Boritsch, E., Vandenbogaert, M., Rodríguez de la Vega, R.C., Thiberge, J.-M., Caro, V., Gaillard, J.-L., Heym, B., Girard-Misguich, F., Brosch, R., Sapriel, G., 2016. Mycobacterial Pan-Genome Analysis Suggests Important Role of Plasmids in the Radiation of Type VII Secretion Systems. Genome Biol. Evol. 8, 387–402. doi:10.1093/gbe/evw001

Elliott, S.R., Tischler, A.D., 2016. Phosphate responsive regulation provides insights for ESX-5 function in Mycobacterium tuberculosis. Curr. Genet. 62, 759–763. doi:10.1007/s00294-016-0604-4

Famelis, N., Rivera-Calzada, A., Degliesposti, G., Wingender, M., Mietrach, N., Skehel, J.M., Fernandez-Leiro, R., Böttcher, B., Schlosser, A., Llorca, O., Geibel, S., 2019. Architecture of the mycobacterial type VII secretion system. Nature 576, 321–325. doi:10.1038/s41586-019-1633-1

Flint, J.L., Kowalski, J.C., Karnati, P.K., Derbyshire, K.M., 2004. The RD1 virulence locus of Mycobacterium tuberculosis regulates DNA transfer in Mycobacterium smegmatis. Proc. Natl. Acad. Sci. USA 101, 12598–12603. doi:10.1073/pnas.0404892101

Fortune, S.M., Jaeger, A., Sarracino, D.A., Chase, M.R., Sassetti, C.M., Sherman, D.R., Bloom, B.R., Rubin, E.J., 2005. Mutually dependent secretion of proteins required for mycobacterial virulence. Proc. Natl. Acad. Sci. USA 102, 10676–10681. doi:10.1073/pnas.0504922102

Garufi, G., Butler, E., Missiakas, D., 2008. ESAT-6-like protein secretion in Bacillus anthracis. J. Bacteriol. 190, 7004–7011. doi:10.1128/JB.00458-08

Gey Van Pittius, N.C., Gamieldien, J., Hide, W., Brown, G.D., Siezen, R.J., Beyers, A.D., 2001. The ESAT-6 gene cluster of Mycobacterium tuberculosis and other high G+C Gram-positive bacteria. Genome Biol. 2, RESEARCH0044. doi:10.1186/gb-2001-2-10-research0044

Glaeser, R.M., Taylor, K.A., 1978. Radiation damage relative to transmission electron microscopy of biological specimens at low temperature: a review. J. Microsc. 112, 127–138.

Gordon, A.H., Hart, P.D., Young, M.R., 1980. Ammonia inhibits phagosome-lysosome fusion in macrophages. Nature 286, 79–80. doi:10.1038/286079a0

Gray, T.A., Clark, R.R., Boucher, N., Lapierre, P., Smith, C., Derbyshire, K.M., 2016. Intercellular communication and conjugation are mediated by ESX secretion systems in mycobacteria. Science 354, 347–350. doi:10.1126/science.aag0828

Gröschel, M.I., Sayes, F., Simeone, R., Majlessi, L., Brosch, R., 2016. ESX secretion systems: mycobacterial evolution to counter host immunity. Nat. Rev. Microbiol. 14, 677–691. doi:10.1038/nrmicro.2016.131

Guerin, E., Cambray, G., Sanchez-Alberola, N., Campoy, S., Erill, I., Da Re, S., Gonzalez-Zorn, B., Barbé, J., Ploy, M.-C., Mazel, D., 2009. The SOS response controls integron recombination. Science 324, 1034. doi:10.1126/science.1172914

Hsu, T., Hingley-Wilson, S.M., Chen, B., Chen, M., Dai, A.Z., Morin, P.M., Marks, C.B., Padiyar, J., Goulding, C., Gingery, M., Eisenberg, D., Russell, R.G., Derrick, S.C., Collins, F.M., Morris, S.L., King, C.H., Jacobs, W.R., 2003. The primary mechanism of attenuation of bacillus Calmette-Guerin is a loss of secreted lytic function required for invasion of lung interstitial tissue. Proc. Natl. Acad. Sci. USA 100, 12420–12425. doi:10.1073/pnas.1635213100

Huppert, L.A., Ramsdell, T.L., Chase, M.R., Sarracino, D.A., Fortune, S.M., Burton, B.M., 2014. The ESX system in Bacillus subtilis mediates protein secretion. PLoS One 9, e96267. doi:10.1371/journal.pone.0096267

Kivisaar, M., 2003. Stationary phase mutagenesis: mechanisms that accelerate adaptation of microbial populations under environmental stress. Environ. Microbiol. 5, 814–827. doi:10.1046/j.1462-2920.2003.00488.x

Landgraf, D., Okumus, B., Chien, P., Baker, T.A., Paulsson, J., 2012. Segregation of molecules at cell division reveals native protein localization. Nat. Methods 9, 480–482. doi:10.1038/nmeth.1955

Leake, M.C., Chandler, J.H., Wadhams, G.H., Bai, F., Berry, R.M., Armitage, J.P., 2006. Stoichiometry and turnover in single, functioning membrane protein complexes. Nature 443, 355–358. doi:10.1038/nature05135

Murphy, K.C., Nelson, S.J., Nambi, S., Papavinasasundaram, K., Baer, C.E., Sassetti, C.M., 2018. ORBIT: a new paradigm for genetic engineering of mycobacterial chromosomes. MBio 9. doi:10.1128/mBio.01467-18

Newton-Foot, M., Warren, R.M., Sampson, S.L., van Helden, P.D., Gey van Pittius, N.C., 2016. The plasmid-mediated evolution of the mycobacterial ESX (Type VII) secretion systems. BMC Evol. Biol. 16, 62. doi:10.1186/s12862-016-0631-2

Pallen, M.J., 2002. The ESAT-6/WXG100 superfamily -- and a new Gram-positive secretion system? Trends Microbiol. 10, 209–212. doi:10.1016/s0966-842x(02)02345-4

Pan, K.Z., Saunders, T.E., Flor-Parra, I., Howard, M., Chang, F., 2014. Cortical regulation of cell size by a sizer cdr2p. Elife 3, e02040. doi:10.7554/eLife.02040

Petridis, M., Benjak, A., Cook, G.M., 2015. Defining the nitrogen regulated transcriptome of Mycobacterium smegmatis using continuous culture. BMC Genomics 16, 821. doi:10.1186/s12864-015-2051-x

Phan, T.H., van Leeuwen, L.M., Kuijl, C., Ummels, R., van Stempvoort, G., Rubio-Canalejas, A., Piersma, S.R., Jiménez, C.R., van der Sar, A.M., Houben, E.N.G., Bitter, W., 2018 EspH is a hypervirulence factor for Mycobacterium marinum and essential for the secretion of the ESX-1 substrates EspE and EspF. PLoS Pathog. 14, e1007247. doi:10.1371/journal.ppat.1007247

Poweleit, N., Czudnochowski, N., Nakagawa, R., Trinidad, D.D., Murphy, K.C., Sassetti, C.M., Rosenberg, O.S., 2019. The structure of the endogenous ESX-3 secretion system. Elife 8. doi:10.7554/eLife.52983

Pym, A.S., Brodin, P., Brosch, R., Huerre, M., Cole, S.T., 2002. Loss of RD1 contributed to the attenuation of the live tuberculosis vaccines Mycobacterium bovis BCG and Mycobacterium microti. Mol. Microbiol. 46, 709–717. doi:10.1046/j.1365-2958.2002.03237.x

Pym, A.S., Brodin, P., Majlessi, L., Brosch, R., Demangel, C., Williams, A., Griffiths, K.E., Marchal, G., Leclerc, C., Cole, S.T., 2003. Recombinant BCG exporting ESAT-6 confers enhanced protection against tuberculosis. Nat. Med. 9, 533–539. doi:10.1038/nm859

Ritchie, M.E., Phipson, B., Wu, D., Hu, Y., Law, C.W., Shi, W., Smyth, G.K., 2015. limma powers differential expression analyses for RNA-sequencing and microarray studies. Nucleic Acids Res. 43, e47. doi:10.1093/nar/gkv007

Sassetti, C.M., Rubin, E.J., 2003. Genetic requirements for mycobacterial survival during infection. Proc. Natl. Acad. Sci. USA 100, 12989–12994. doi:10.1073/pnas.2134250100

Serafini, A., Boldrin, F., Palù, G., Manganelli, R., 2009. Characterization of a Mycobacterium tuberculosis ESX-3 conditional mutant: essentiality and rescue by iron and zinc. J. Bacteriol. 191, 6340–6344. doi:10.1128/JB.00756-09

Serafini, A., Pisu, D., Palù, G., Rodriguez, G.M., Manganelli, R., 2013. The ESX-3 secretion system is necessary for iron and zinc homeostasis in Mycobacterium tuberculosis. PLoS One 8, e78351. doi:10.1371/journal.pone.0078351

Siegrist, M.S., Steigedal, M., Ahmad, R., Mehra, A., Dragset, M.S., Schuster, B.M., Philips, J.A., Carr, S.A., Rubin, E.J., 2014. Mycobacterial Esx-3 requires multiple components for iron acquisition. MBio 5, e01073–14. doi:10.1128/mBio.01073-14

Siegrist, M.S., Unnikrishnan, M., McConnell, M.J., Borowsky, M., Cheng, T.-Y., Siddiqi, N., Fortune, S.M., Moody, D.B., Rubin, E.J., 2009. Mycobacterial Esx-3 is required for mycobactin-mediated iron acquisition. Proc. Natl. Acad. Sci. USA 106, 18792–18797. doi:10.1073/pnas.0900589106

Smyth, G.K., 2004. Linear models and empirical bayes methods for assessing differential expression in microarray experiments. Stat Appl Genet Mol Biol 3, Article3. doi:10.2202/1544-6115.1027

Soler-Arnedo, P., Sala, C., Zhang, M., Cole, S.T., Piton, J., 2020. Polarly Localized EccE1 Is Required for ESX-1 Function and Stabilization of ESX-1 Membrane Proteins in Mycobacterium tuberculosis. J. Bacteriol. 202. doi:10.1128/JB.00662-19

Sørensen, A.L., Nagai, S., Houen, G., Andersen, P., Andersen, A.B., 1995. Purification and characterization of a low-molecular-mass T-cell antigen secreted by Mycobacterium tuberculosis. Infect. Immun. 63, 1710–1717. doi:10.1128/IAI.63.5.1710-1717.1995

Stanley, S.A., Raghavan, S., Hwang, W.W., Cox, J.S., 2003. Acute infection and macrophage subversion by Mycobacterium tuberculosis require a specialized secretion system. Proc. Natl. Acad. Sci. U. S. A. 100, 13001–13006.

Tufariello, J.M., Chapman, J.R., Kerantzas, C.A., Wong, K.-W., Vilchèze, C., Jones, C.M., Cole, L.E., Tinaztepe, E., Thompson, V., Fenyö, D., Niederweis, M., Ueberheide, B., Philips, J.A., Jacobs, W.R., 2016. Separable roles for Mycobacterium tuberculosis ESX-3 effectors in iron acquisition and virulence. Proc. Natl. Acad. Sci. USA 113, E348–57. doi:10.1073/pnas.1523321113

Vince, A., Dawson, A.M., Park, N., O’Grady, F., 1973. Ammonia production by intestinal bacteria. Gut 14, 171–177. doi:10.1136/gut.14.3.171

Way, S.S., Wilson, C.B., 2005. The Mycobacterium tuberculosis ESAT-6 homologue in Listeria monocytogenes is dispensable for growth in vitro and in vivo. Infect. Immun. 73, 6151– 6153. doi:10.1128/IAI.73.9.6151-6153.2005

Williams, K.J., Bryant, W.A., Jenkins, V.A., Barton, G.R., Witney, A.A., Pinney, J.W., Robertson, B.D., 2013. Deciphering the response of Mycobacterium smegmatis to nitrogen stress using bipartite active modules. BMC Genomics 14, 436. doi:10.1186/1471-2164-14-436

Wirth, S.E., Krywy, J.A., Aldridge, B.B., Fortune, S.M., Fernandez-Suarez, M., Gray, T.A., Derbyshire, K.M., 2012. Polar assembly and scaffolding proteins of the virulence-associated ESX-1 secretory apparatus in mycobacteria. Mol. Microbiol. 83, 654–664. doi:10.1111/j.1365-2958.2011.07958.x

